# Chloroform-injection (CI) and Spontaneous-phase-transition (SPT) are Novel Methods, Simplifying the Fabrication Liposomes

**DOI:** 10.1101/2020.08.09.242693

**Authors:** Muhammad Ijaz Khan, Naveed Ahmed, Muhammad Farooq Umer, Amina Riaz, Nasir Mehmood Ahmad, Gul Majid Khan

## Abstract

Intricate formulation methods and/or use of sophisticated equipment limit the prevalence of liposomal dosage-forms. Simple techniques are developed to assemble amphiphiles into globular lamellae while transiting from immiscible organic to the aqueous phase. Various parameters are optimized by injecting chloroform solution of amphiphiles into the aqueous phase and subsequent removal of the organic phase. Further simplification is achieved by reorienting amphiphiles through a spontaneous phase transition in a swirling biphasic system during evaporation of the organic phase under vacuum. Although the chloroform injection yields smaller size and PDI yet spontaneous phase transition method overrides simplicity and productivity. The size distribution of liposomes and solid/solvent ratio in both or any phases of formulation show direct relation. Surface charge dependant large unilamellar vesicles with a narrow distribution have PDI <0.4 in 10 μM saline. As small and monodisperse liposomes are prerequisites in targeted drug delivery strategies. Hence the desired size distribution <200 d.nm and PDI <0.15 is obtained through serial membrane-filtration method. Phosphatidylcholine/water 4 μmol/ml is achieved at a temperature of 10°C below the phase-transition temperature of phospholipids ensuing suitability for thermolabile entities and high entrapment efficiency. Both methods furnish the de-novo rearrangement of amphophiles into globular lamellae aiding in the larger entrapped volume. The immiscible organic phase facilitates faster and complete removable of the organic phase. High cholesterol content (55.6 mol%) imparts stability in primary hydration medium at 5+3°C for 6 months in light-protected type-1 glass vial. Collectively the reported methods are novel, scalable, time-efficient yielding high productivity in simple equipment.

## 1 Introduction

Liposomes are globular vesicles of amphiphiles having aqueous core concealed in lipophilic lamellae. Phospholipids coalesce in a manner that hydrophilic heads hold the water on both sides while hydrophobic-tails rootle away in the bilayers. The dual-compartment structure enables them to entrap both hydrophilic and lipophilic drugs in the core and the bilayer respectively [1, [2]. Liposomal classification is based on morphology (uni/multilamellar), dimensions (giant, large or small), surface-charge (anionic, cationic or neutral) and function (conventional, coated, targeted, triggered, stealth, slow-release or combinatorial) [3]. Liposomal dosage-forms enhance the stability and or biodistribution of entrapped drugs [4, [5]. However, the quality elements including size, morphology, surface charge, and ligand if attached essentially affect the pharmacokinetic behaviour of entrapped drugs [6, [7]. Resemblance with bio-membranes, [8] enhanced permeability and retention effect [9] and or active targeting [10, [11, [12] confer them improved pharmacological profile. Consequently, the liposomal drugs show enhanced therapeutic efficacy and safety [13, [14] than conventional dosage forms. Therefore, liposomes are extensively employed in therapeutic and theranostic targeted drug delivery systems [15, [16].

The structural resemblance with biological membranes upheld the liposomes as tempting biocompatible drug carriers [8]. However, the knotty formulation techniques and or sophisticated equipment hampered their prevalence. Consequently besides a variety of reported methods, scaleup to industrial manufacturing and scale down for personalized treatment remained challenging [17]. That’s why the film hydration method (Bingham et al; 1965) with or without modifications is still accustomed due to simple procedure and equipment [2, [4, [15]. Organic solvent is either removed before hydration with aqueous phase (film hydration methods) or from the mixture of both phases (bulk methods) [3]. The rest is manipulation of these basic principles for good results. Bulk methods include reverse-phase evaporation, [18] Organic-phase injection, [19, [20, [21, [22] solvent spherules formation, [23] rapid preparation of giant liposomes [24] detergent dialysis, [25, [26] rapid solvent exchange, [27] microfluidics, [28, [29] use of supercritical fluids [30] and modified electro formation [31]. Majority of these methods require sophisticated equipment and/or convoluted procedures that confine the commonness of the dosage-form.

Film hydration methods rout the film forming components through intermediary solid phase [2, [4]. Therefor subsequent film hydration usually takes much longer and result in compositional inhomogeneity, particularly in the presence of cholesterol (Cho) [27]. Bulk methods for preparation of liposomes are preferred for their ease, lamellar homogeneity and good entrapment efficiency. However organic phase plays a pivotal role in such methods for consigning phospholipids into the aqueous phase. Some of the bulk methods used water miscible organic solvents [20, [24] which are difficult to be removed from the final formulation. The residues of such miscible solvents impede the stability of vesicles, denature the susceptible drugs and also toxic to human health. Cho is essential component to impart rigidity and hence physical stability to liposomes [32, [33]. The absence of Cho in some of bulk methods undoubtedly facilitated vesiculation but resulted in delicate vesicles and hence prone to leakiness in shelf life storage.

Simple methods are reported here for instant preparation of liposomes with immiscible organic solvent, high Cho contents and bypassing intermediary solid film assuring lamellar homogeneity [27]. Customarily the methods were titled as chloroform-injection (CI) and spontaneous-phase-transition (SPT) methods. CI method comprised the injection of amphiphiles and subsequent removal of immiscible organic-phase. While in single-step SPT method evaporation under reduced pressure from the biphasic mixture induced a de-novo reorientation of amphiphiles into lamellar vesicle. Complete removal of organic phase was achieved due to its immiscibility, low boiling point and evaporation under reduced pressure. The resultant unilamellar vesicles were sized through serial-membrane filtration [34, [35] to obtain the desired size ˂200 nm and PDI ˂0.15. Larger entrapped volume was attributed to the de-novo rearrangement of amphiphiles as in contact with the aqueous phase. Although CI-method yields a comparatively smaller size and narrower distribution but more handling losses than the SPT-method. High solid/solvents ratio, lower temperature, short processing time, scalability and simple procedure and equipment are distinctive features of the reported methods.

## 2 Material and Methods

Hydrogenated phosphatidylcholine egg yolk, (HPCE); was gifted by Lipoid GMBH, Frigenstrass-4, Ludwigshafen, Germany. Cholesterol (Cho), Rhodamine-B (RhB), Chloroform analytical grade (CHCl_3_) were obtained from Sigma Aldrich. Nylon membrane filters 0.4 μ and 0.2 μ (Sartorius, Germany), Dialysis membrane (12-14 kDa) (Membrane Filtration Products-Texas, USA). Syringes with a needle of 0.16 d.mm (bore-dia) were purchased and blunted their tips with sandpaper. Freshly prepared double-distilled water (DW) was duly filtered with 0.1 μm membrane-filter prior to use. Stock solutions of HPCE/CHCL_3_ 1.00 mmol/ml and Cho/CHCL_3_ 1.00 mmol/ml were stored at 5±3°C.

### 2.1 Chloroform injection method

The required quantity of HPCE/Cho/CHCl_3_ was filled in the syringe with a 0.16 d.mm blunt needle. Aqueous phase was adjusted to prerequisite temperature (25, 35, 45 or 55°C) at high stirring speed but vortexing was avoided to prevent accumulation of solids at the base of the device. Organic phase was promptly injected into just below the surface of the aqueous phase. The mixture was allowed to mix at 10 rpm for next 2-3 min. A milky suspension obtained was transferred to the rotary evaporator and allowed to evaporate under reduced pressure (through a 16-psi vacuum pump) at ∟45°, prerequisite temperature and 150 rpm for 20 min. A maximum of half of the rotating flask was occupied to allow sufficient surface area for evaporation.

During optimization studies, unreacted material and or giant vesicles were removed through slow centrifugation at 25°C, 1000 g for 3 min with zero acceleration deacceleration. However, optimized formulations were sized through serial membrane filtration immediately while maintaining the process temperature. Replicate trials were produced to ensure the reproducibility and quality of vesicles. Various formulation and process parameters were optimized including variable quantities of organic phase, aqueous phase, Cho concentration and temperature while the quantity of phospholipids was kept constant.

### 2.2 Liposome preparation by SPT method

Chronologically to improve efficiency in terms of processing time, yield and or material loss the optimized parameters of CI-method were further investigated through the SPT-method. DW was brought to required temperature in the rotating flask of a rotary evaporator. HPCE/Cho/CHCl_3_ was gently added, partitioning at bottom of the flask. The mixture was allowed to rotate at ∟45°, 150 rpm for 20 minutes under reduced pressure at 45°C to evaporate the immiscible organic phase. Liposomes were produced de-novo by spontaneous reorientation of amphiphiles while transiting from organic to aqueous phase. The vesicles produced by SPT-method were evaluated and compared with CI-method under a given set of conditions. Highly concentrated LUVs HPCE/DW 4 μmol/ml were sized through serial membrane filtration at the processing temperature.

### 2.3 Sizing and purification

Sizing of liposomes was obligatory to obtain the desired size and narrow distribution by serial membrane filtration. [34, [35] Briefly, the freshly prepared liposomes were passed 5x each through 0.4 μm followed by a 0.2 μm nylon membrane filter. Sizing was performed in the primary hydration medium at the optimized processing temperature.

The sized vesicles were purified through membrane dialysis to remove non-liposomal contents if required. Concisely, the sized vesicles in primary hydration volume were transferred to the dialysis sac (12-14 kDa MW cut-off) and placed in 10x freshly prepared distilled water. Temperature was maintained at 5±3°C and stirring at 10 rpm for 24 hrs. Washing media was replaced every 6hrs to ensure equilibration. Product was collected and stored in type-1 glass vials duly wrapped in aluminium foil at 5±3°C till further use

### 2.4 Dynamic light scattering

The formulations were analysed for Z-average hydrodynamic diameter (Z-av), poly-dispersity-index (PDI) and zeta potential (ζ-potential) using dynamic light scattering (DLS) technique through Zeta-sizer ZS90 (Malvern Instruments, Malvern, UK). Size analysis was carried out at 25°C, scattering-angle 90° in aqueous dispersion at 25°C having RI 1.33, viscosity 1 cps and dielectric constant 79. To prevent multi-scattering, 100x dilution in DW at 25°C was used for comparative size distribution. Subsequently, to minimize the effect of surface potential, 100x dilution of liposomal stock dispersion in 10 μM NaCl (clarified through o.2μ membrane filter) was also analysed for comparative size distribution.

### 2.5 Scanning electron microscopy

Scanning electron microscope (SEM) JSM 6490A, JEOL, Japan was used to study and compare the size and morphology of liposomes. Unsized samples produced by CI and SPT methods under the same set of optimized parameters (Section-2.1, 2.2) were subjected to comparative analysis. 50 μl from 100x diluted stock dispersion was dropped on the silicon wafers and gently blot dried after 2-3 minutes. Surface water was evaporated at room temperature for 5 minutes. The dried smear was gold-coated in Joel 1100 (JFC Ion Sputter) at 50 mA in argon environment (50Pa) for 50 seconds. Coated samples were observed at 10,000x magnification under the SEM mod.

### 2.5 Atomic force microscopy

The topography of optimized formulations after sizing was performed with a Scanning Probe Microscope (SPM) JSPM-5200, JEOL, Japan. Imaging was performed at 20-25°C in DW on 1cm^2^ silicon wafers. 50 μl of 100x diluted stock dispersion was dropped on a silicon wafer to minimize clustering and sticking of the vesicles. The samples were gently blotted after 2 mins with filter paper. Excessive water from the smear was evaporated at room temperature before loading the sample on specimen stage. All samples were observed within 5-15 min of deposition. The AFM was operated in amplitude-detection (AC-AFM) mode in atmospheric conditions. Aluminium back-coated silicon nitride tip was used for scanning. AC-AFM scans the sample by vibrating cantilever with constant amplitude in an area of a specific frequency at distance 5-20Å. [36] The change in the voltage (error signal) for a constant amplitude of cantilever is computed into an image. The cantilever was moved to approach the sample to a point where inter-atomic forces interact to scan the sample and confirmed through reference voltage and Z-piezo position. Images were scanned and analysed in scanning and processing mode respectively through WINSPM 5 software. Topographical images along with error signals were obtained showing minor superficial variation in the samples. The diameter and height of the vesicles was measured by drawing a line with pointers across the images. Surface roughness was directly estimated from the arithmetic average roughness (Ra) values in height mode. The volume of vesicles was calculated by using the following formula;

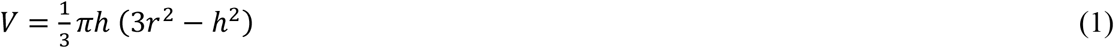

Where ‘r’ represents the radius and ‘h’ height of the vesicles.

The rigidity of liposomes is the measure of resistance to deformation in a set of given circumstances. In-vitro rigidity of liposomes was calculated from the height data of AC-AFM and size data from DLS using 10 μM NaCl as a dilution medium to compress the surface charge cloud. [37] Average rigidity of the samples was calculated through following equation.

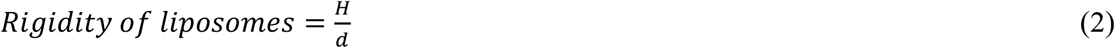

Where ‘H’ is the height from AFM data and ‘d’ is the diameter of vesicles from DLS data.

### 2.6 Entrapped volume

Aqueous volume entrapped by liposomes is indicative of the process efficiency and consequent encapsulation of dissolved drugs. RhB was dissolved in DW at a concentration of 500 μmol/ml. This solution was used to prepare triplicate samples of liposomes at variable concentrations of HPCE/CHCl_3_ in a constant volume of the aqueous phase. Samples were prepared by using optimized conditions (Section-2.2). The RhB-liposomes were sized through serial membrane filtration and purified by exhaustive dialysis (Section-2.3). 100 μl of dialyzed liposomes were added to 3 ml of DW containing 2% Triton X-100 to avoid liposomal turbidity and relieve the self-quenched RhB. The mixture was transferred to a quartz cuvette and absorbance was measured at λ-max 550 nm while taking DW as a blank. RhB solution without phospholipids was analysed to calculate any traces of unentrapped RhB. The observed values were quantitated against a standard curve of RhB solution in DW. Each sample was also analysed for phospholipid concentration by phosphate analysis. Entrapped volume was calculated by;

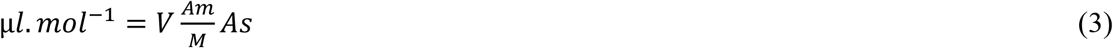

where ‘M’ represents a mole of phospholipids in 3 ml dialyzed liposomes, ‘Am’ is the measured absorbance and ‘As’ denotes standard absorbance in known volume V (*L*) [22].

### 2.7 Accelerated stability studies

ICHQ protocol Q1A(R2)2.2.7.4 was followed to test the accelerated stability of liposomes intended to be stored in refrigeration [38]. The liposomal preparation containing HPCE/Cho as 4:5 molar ratio and HPCE/DW 4 μmol/ml (Section-2.2, 2.3) were exposed to accelerated storage conditions. The liquid samples in primary hydration media were stored in Type-1 glass vials aptly wrapped in aluminium foil and each kept at RT, 23±2°C and REF, 5± 3°C for 6 months. Being stored in impermeable and light-resistant containers only thermal stability was a concern of probable instability. Additional test points were included to observe any change in a small interval of time including day 0, 30, 60, 90 and 180. Physical appearance in terms of uniformity or precipitation/aggregation was considered decisive to the stability of various samples. Samples at a given interval were analysed for Z-av and PDI (Section-2.4).

### 2.8 Data reporting

All experiments were performed in replicate with a variable n value. Optimization experiments were reproduced in duplicate while optimized formulations were reproduced multiple times to ensure reproducibility and as required to study various parameters. Data was averaged and presented as ± standard deviation.

## 3 Results

### 3.1 Key outcomes of optimization studies

The samples were assessed on the basis of Z-av, PDI and physical uniformity during formulation. In the same group of experiments count-rate (KCP) during DLS measurement was considered as a measure of comparative vesiculation. Smaller size, PDI and higher count rate at given sample dilution were considered for shortlisting.

#### 3.1.1 Optimization of the organic phase

Liposomes were prepared through CI-method (Section-2.1) at 25°C and hydrogenated phosphatidyl choline from egg yolk/double distilled water (HPCE/DW) 0.15 μmol/ml. Z-av was slightly declining within the same set of experiments by diluting phospholipids in the organic-phase (Table-1a) phosphatidyl choline from egg yolk/chloroform (HPCE/CHCl_3_) 8 - 2 μmol/ml. Further dilution yet again resulted in a slight increase in size. However, both PDI and KCPs were continuously improving with increasing dilutions of HPCE/CHCl_3_. Another set of experiments (Table-1d) was performed using HPCE/DW 0.38 μmol/ml and variable HPCE/CHCl_3_ to evaluate the minimum required volume of organic solvent. All the above-mentioned parameters were increasing with increasing concentration of HPCE/CHCl_3_ from 10 to 30 μmol/ml. While precipitation was observed at HPCE/CHCl_3_ 60 μmol/ml.

**Table 1.**
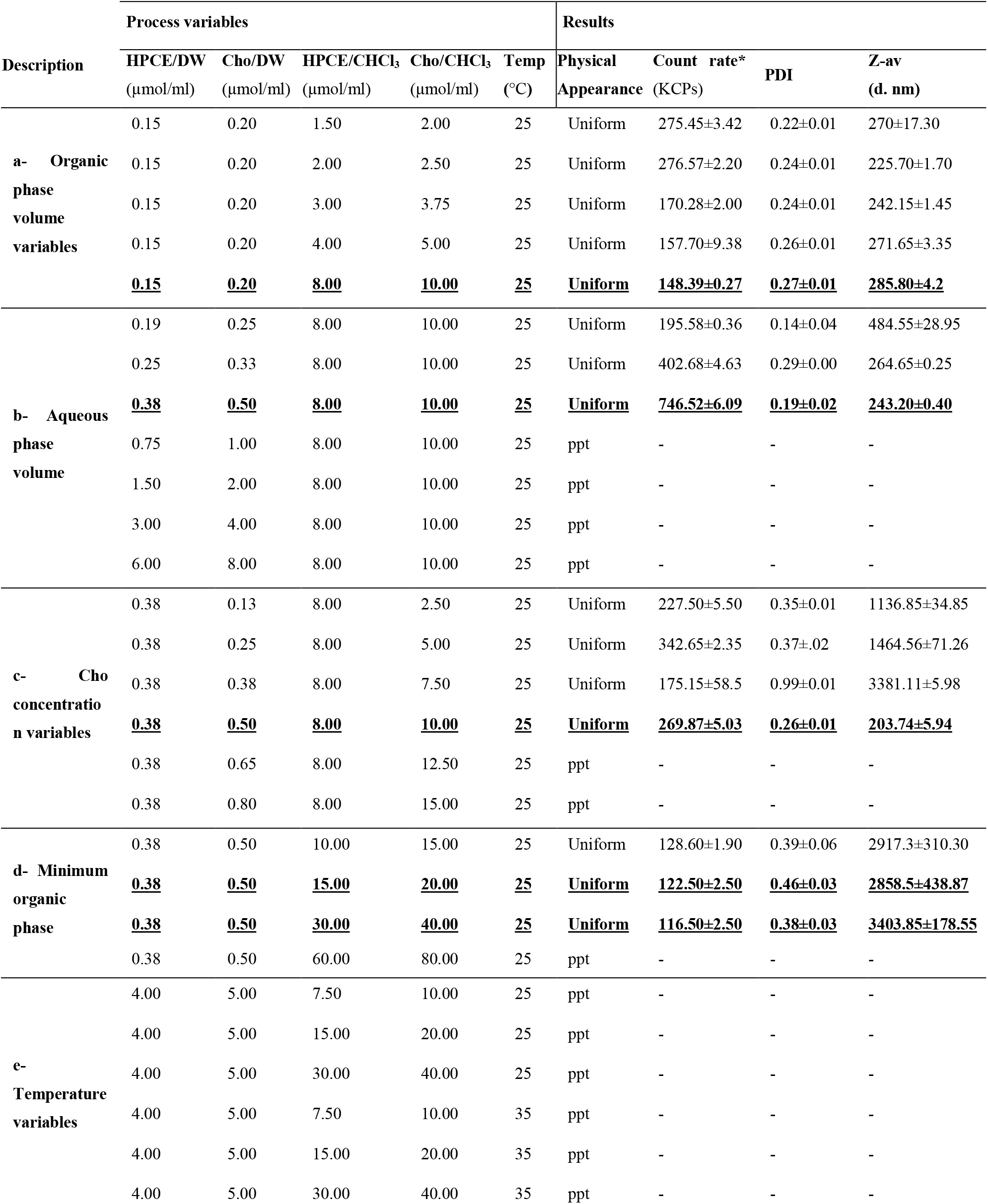
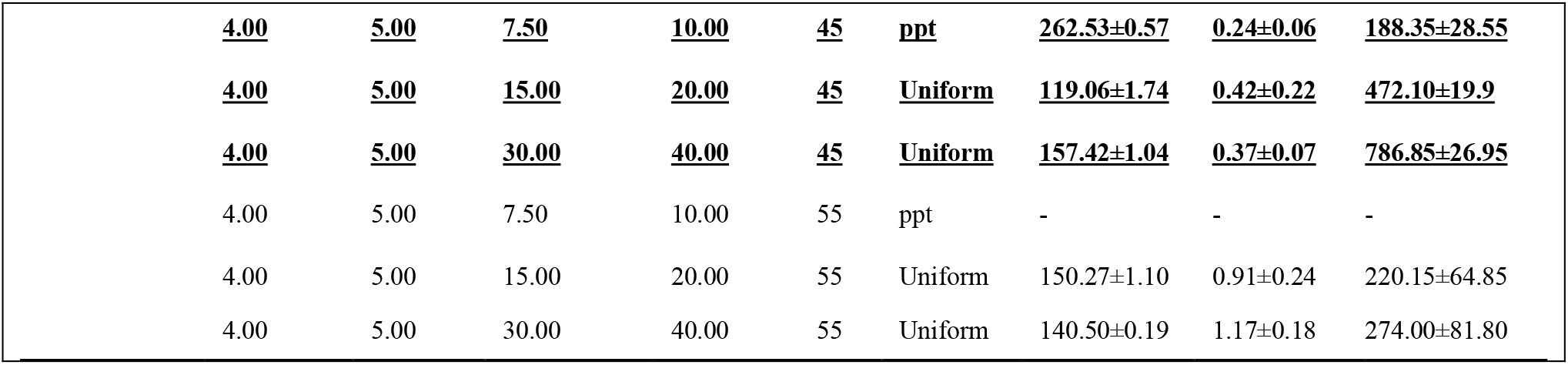
Key results of optimization studies through CI-method. The shortlisted results were highlighted and considered as a base for the next experiment ^**a:**^ Quantity of organic-phase was varied against a fixed volume of aqueous-phase ^**b:**^ The quantity of aqueous-phase was reduced against the pre-set quantity of organic-phase from step-a ^**c:**^ Quantity of maximum Cho concentration against the pre-set quantity of HPCE was assessed with the optimized parameters of a & b. ^**d:**^ Minimum volume of organic-phase required for vesiculation was assessed with pre-set a, b & c. ^**e:**^ Maximum concentration of amphiphiles in minimum solvent was assessed through variable temperature to evaluate minimum possible temperature for vesiculation **\***Size measurement was performed at standard dilution for a particular set of experiments.

#### 3.1.2 Optimization of aqueous phase

CI-method at 25°C, HPCE/CHCl_3_ 8 μmol/ml and variable aqueous volume were used. The vesicle size was reduced with increasing concentration of solids in the aqueous phase (Table-1b) until HPCE/DW 0.38 μmol/ml. While KCPs were markedly increasing towards concentrating aqueous suspensions. The CI-method at 25°C was effective for dilute HPCE/DW ≤0.38 μmol/ml. Further increment of solids resulted in precipitation and sticking of material to the walls of the rotating flask.

#### 3.1.3 Optimization of HPCE/Cho molar ratio

The liposomes were prepared by CI-method at 25°C, HPCE/DW 0.38 μmol/ml and HPCE/CHCl_3_ 8 μmol/ml. HPCE/Cho ~4:5 molar ratio exhibited better results in terms of uniformity, smaller size, and PDI (Table-1c) without treatment for sizing. The lower molar concentration of Cho showed broader distribution and hydrodynamic diameter while a higher ratio exhibited precipitation against the pre-set quantity of HPCE.

#### 3.1.4 Optimization of processing temperature

Liposomes were formulated at variable temperatures of 25, 35, 45 and 55°C by CI-method with HPCE/Cho 4:5 molar ratio and high solids/solvent ratio (Table-1e). Preoptimized concentration of HPCE/DW 4 μmol/ml, HPCE/CHCl_3_ 7.5, 15 or 30 μmol/ml were evaluated at above-mentioned temperatures. Precipitation was observed at 25°C and 35°C while at 55°C aggregation was observed with diluted HPCE/CHCl_3_ 7.5 μmol/ml. Uniform suspension of vesicles with higher KCP was obtained at 45°C i.e. below the transition temperature of lipid employed. Serial dilutions of HPCE/CHCl_3_ was directly related to smaller size vesicles at any given temperature and volume of the aqueous phase.

### 3.2 Parameters of optimised formulation

The optimized formulations comprised HPCE/Cho 4:5 molar ratio, HPCE/DW 4 μmol/ml, HPCE/CHCl_3_ 7.5, 15 or 30 μmol/ml (variable concentration) or DW/CHCl_3_ 16:1, 4:1 and 2:1 v/v. Optimized conditions included 45°C temperature followed by rotary evaporation under reduced pressure (using 16 psi vacuum pump) at ∟45° and 150 rpm. Different protocols for SPT and CI methods already explained in Sections-2.1 and 2.2 respectively. The liposomes produced by both methods were sized and purified (when required) mentioned in Section-2.3. Optimized parameters were decided on the basis of lower PDI, Z-av, physical uniformity and high solid to solvent ratio at a possible lower temperature as mentioned in Sections-3.1.1, 3.1.2, 3.1.3 and 3.1.4. Various problems encountered during experimentation were effectively resolved and summarized in Table-2.

**Table 2.**
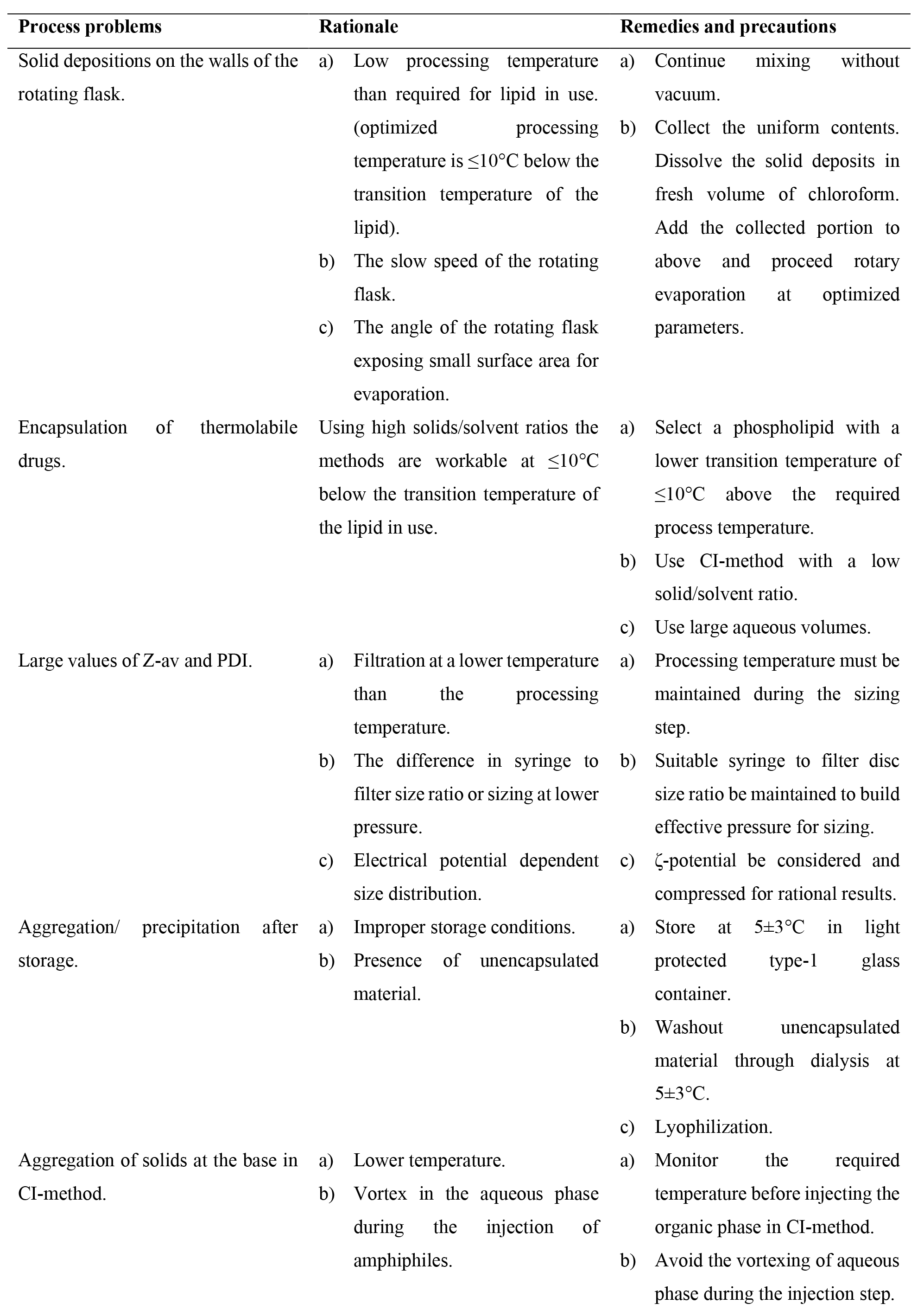
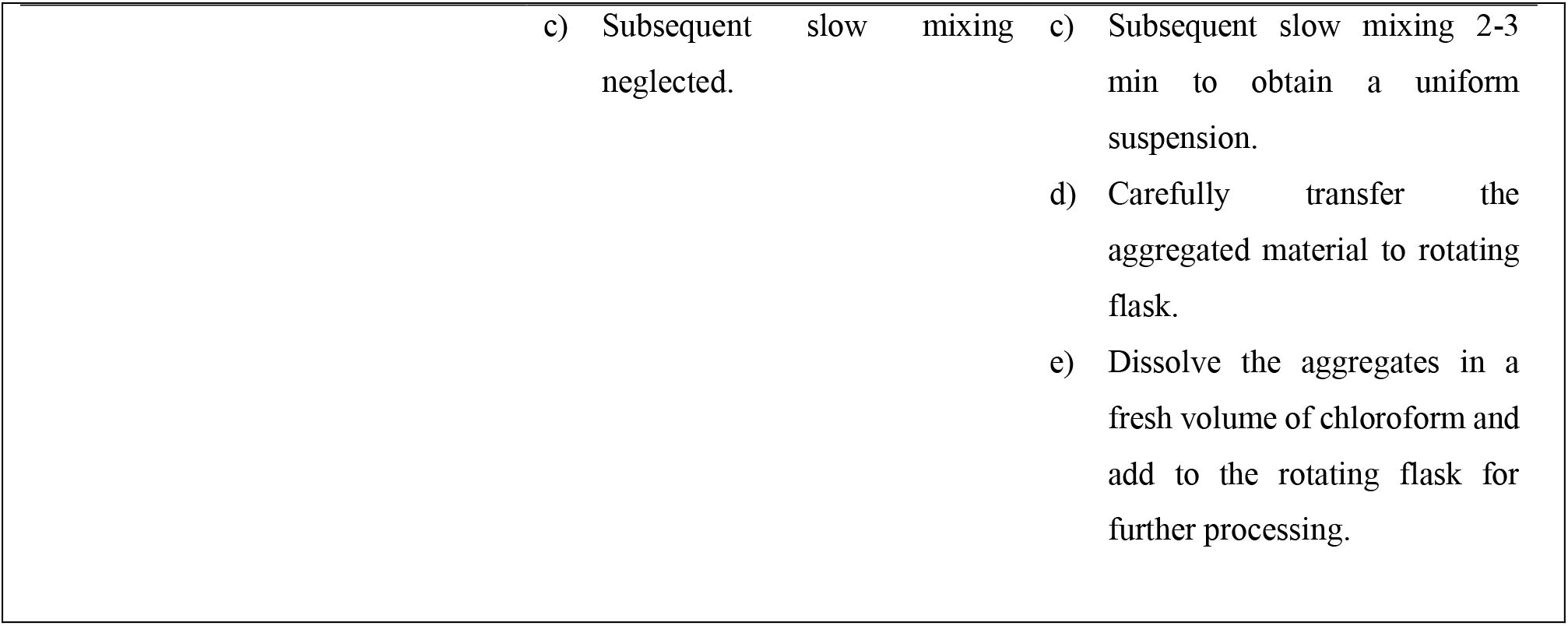
Troubleshooting of expected processing problems while preparing liposomes with CI and SPT methods.

| Process problems | Rationale | Remedies and precautions |
| --- | --- | --- |
| Solid depositions on the walls of the rotating flask. | a) Low processing temperature than required for lipid in use. (optimized processing temperature is $\leq 10^{\circ}\text{C}$ below the transition temperature of the lipid).<br>b) The slow speed of the rotating flask.<br>c) The angle of the rotating flask exposing small surface area for evaporation. | a) Continue mixing without vacuum.<br>b) Collect the uniform contents. Dissolve the solid deposits in fresh volume of chloroform. Add the collected portion to above and proceed rotary evaporation at optimized parameters. |
| Encapsulation of thermolabile drugs. | Using high solids/solvent ratios the methods are workable at $\leq 10^{\circ}\text{C}$ below the transition temperature of the lipid in use. | a) Select a phospholipid with a lower transition temperature of $\leq 10^{\circ}\text{C}$ above the required process temperature.<br>b) Use CI-method with a low solid/solvent ratio.<br>c) Use large aqueous volumes. |
| Large values of Z-av and PDI. | a) Filtration at a lower temperature than the processing temperature.<br>b) The difference in syringe to filter size ratio or sizing at lower pressure.<br>c) Electrical potential dependent size distribution. | a) Processing temperature must be maintained during the sizing step.<br>b) Suitable syringe to filter disc size ratio be maintained to build effective pressure for sizing.<br>c) ζ-potential be considered and compressed for rational results. |
| Aggregation/ precipitation after storage. | a) Improper storage conditions.<br>b) Presence of unencapsulated material. | a) Store at $5 \pm 3^{\circ}\text{C}$ in light protected type-1 glass container.<br>b) Washout unencapsulated material through dialysis at $5 \pm 3^{\circ}\text{C}$.<br>c) Lyophilization. |
| Aggregation of solids at the base in CI-method. | a) Lower temperature.<br>b) Vortex in the aqueous phase during the injection of amphiphiles. | a) Monitor the required temperature before injecting the organic phase in CI-method.<br>b) Avoid the vortexing of aqueous phase during the injection step. |

|  |  |
| --- | --- |
| c) Subsequent slow mixing neglected. | c) Subsequent slow mixing 2-3 min to obtain a uniform suspension. |
|  | d) Carefully transfer the aggregated material to rotating flask. |
|  | e) Dissolve the aggregates in a fresh volume of chloroform and add to the rotating flask for further processing. |

### 3.3 Comparison of CI and SPT methods

For comparative studies, vesicles were produced through both CI and SPT methods (Sections-2.1, 2.2). Optimized parameters were used with 2x increasing concentration of HPCE/CHCl_3_ ≥7.5 μmol/ml in organic-phase against pre-set quantity of HPCE/DW 4 μmol/ml (Section 3.2). SPT-method was a further simplified form of CI-method bypassing the cumbersome organic phase injection (Figur-1a & b) and subjecting the whole material for vesiculation.

**Figure 1.**
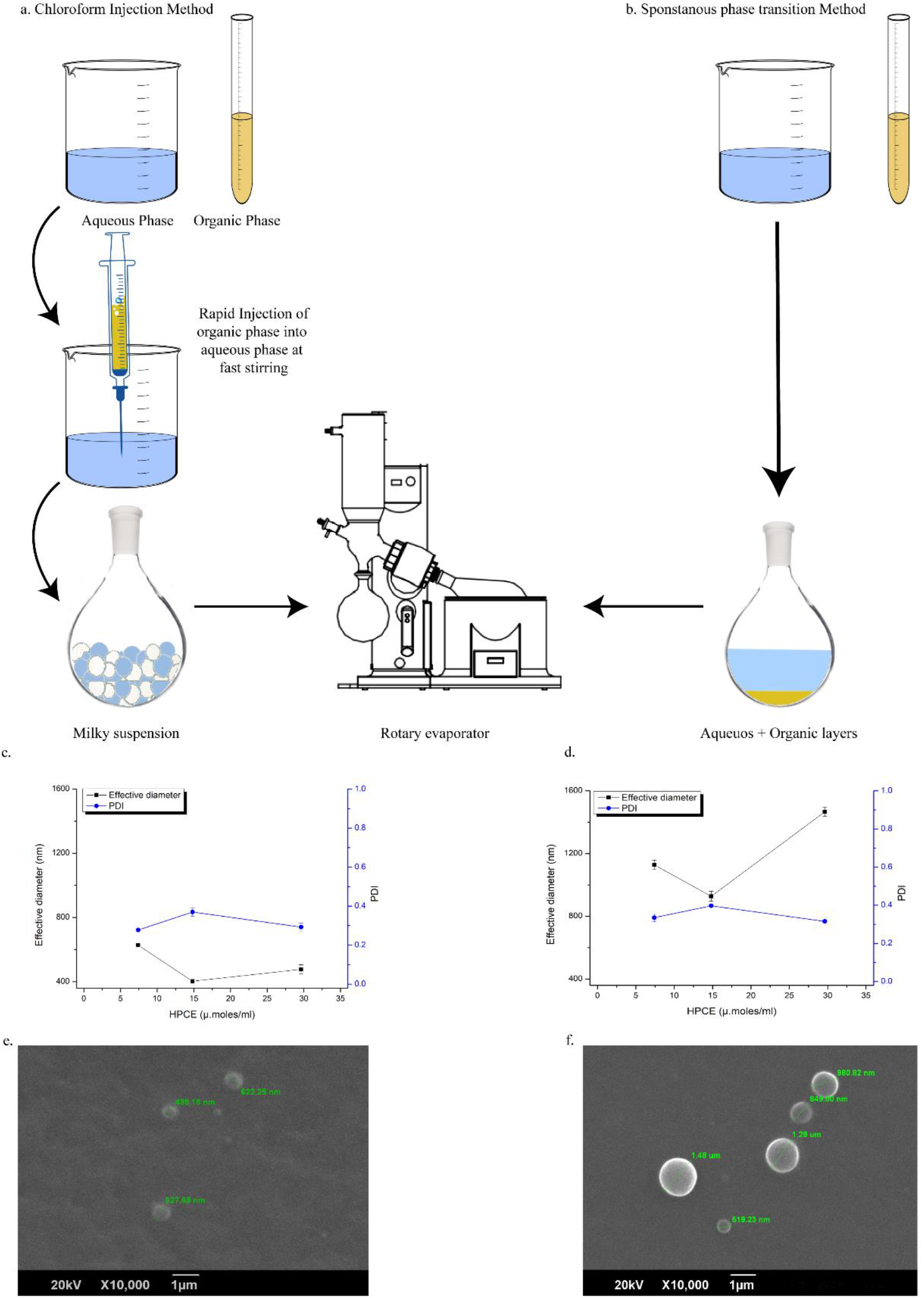
Comparison of CI and SPT liposomes produced with optimized parameters and without sizing ^**a:**^ CI method ^**b:**^ SPT method ^**c:**^ Z-av and PDI of unsized CI-liposomes ^**d:**^ Z-av, and PDI of unsized SPT-liposomes ^**e:**^ SEM image of unsized CI-liposomes ^**f:**^ SEM image of unsized SPT-liposomes

#### 3.3.1 Size distribution analysis

Comparable results in terms of size distribution and PDI of the vesicles were obtained through both CI and SPT methods. Slightly larger Z-av and PDI values were observed through SPT than CI method (Figure-1c, d, e & f) under the same set of optimized parameters. The formulations produced with the SPT-method were more turbid and gave higher KCPs over 100x dilution than the CI-method. Low solids/solvent ratio in the organic-phase resulted in a smaller Z-av and PDI values of the vesicles against a pre-set HPCE/DW 4μmol/ml. The results showed a parabolic pattern with the smallest size at DW/CHCl_3_ 1:4 or HPCE/CHCl_3_ 15μmol/ml. While slight variation in PDI was observed within the same group of experiments.

#### 3.3.2 Effect of sizing by serial filtration

Size reduction by membrane filtration (Section-2.3) resulted in low PDI and Z-av values (Table-3). The size of filtered vesicles was consistent with the pore size of the membrane used for filtration. The lower count-rate for the same set of experiments with 100x dilution indicated a loss in the number of vesicles per ml of the aqueous phase. Since sizing down by membrane filtration was necessary for desired size and PDI, the SPT method was preferred for better yield, processing time, convenience and hence cost-effectiveness.

**Table 3.**
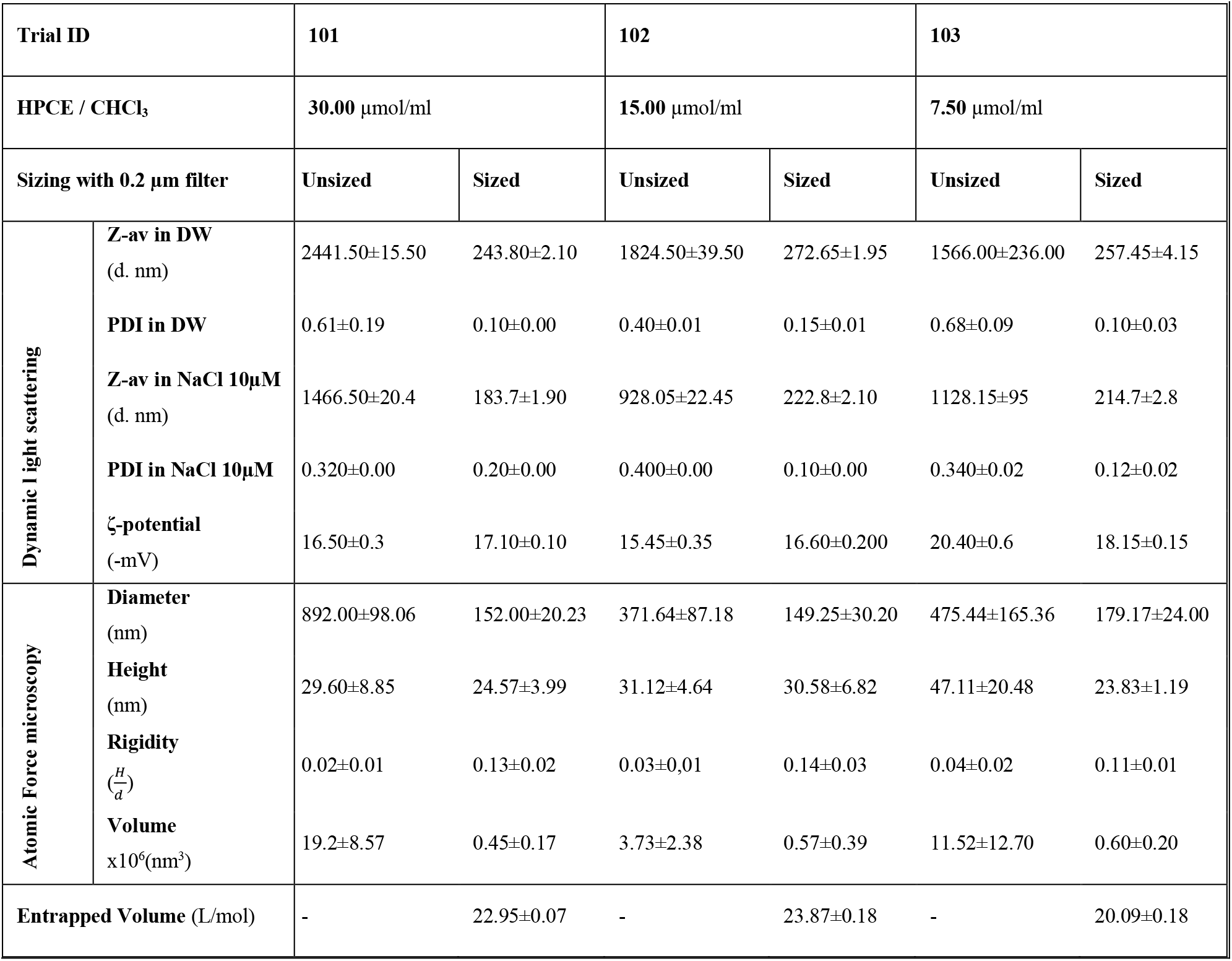
Comparative study of optimized liposomes produced by SPT method at a variable concentration of organic phase before and after sizing with serial membrane filtration

#### 3.3.3 Scanning Electron Microscopy

The morphological analysis under SEM revealed a circular appearance with smooth surfaces of liposomes produced by both CI and SPT methods (Figure-1d & e). A sharp clear outer boundary without any concentric layers confirmed a unilamellar structure covered by the surface membrane. However, SEM images showed an obvious difference in size amongst CI and SPT liposomes without sizing. SPT liposomes were larger in size and comparatively having broader distribution 1023.18±336.44 d.nm. While CI liposomes showed 581.71±61.22 d.nm were comparatively smaller in size and more uniform than SPT liposomes.

### 3.4 Properties of optimized liposomes

Both CI (Section-2.1) and SPT (Section-2.2) methods under optimized conditions (Section-3.2) yielded LUVs. The LUVs were sized by serial membrane filtration (Section-2.3) for narrow size distribution. Sized liposomes were consistent with the smallest pore size of the membrane used, lower PDI and size-dependent variable ζ-potential (Table-3). After careful handling, a 16.7% loss was observed to the initial volume of aqueous phase used. Half of the loss was attributed to the evaporation of water under reduced pressure signifying the complete removal of the organic phase.

#### 3.4.1 Size distribution and surface charge analysis

SPT method under optimized conditions (Section-3.2) yielded LUVs with Z-av ≤ 1466±28.84 d.nm PDI ≤ 0.39±0.003 dependent on varying concentrations of HPCE/CHCl_3_. CI method also furnished LUVs but comparatively of smaller size ≤627.4±0.99 d.nm and PDI ≤0.37±0.020 (Figure-1c &d). More handling losses were observed due to the injection step of the CI-method (Figure-1a, b). DLS analysis in DW and 10 μM NaCl showed different but comparable results indicating ζ-potential dependent size variation (Table-3). The Z-av of unsized liposomes was declining with further dilution of solids in an organic solvent for a pre-set quantity of the aqueous phase (Table-3). PDI and ζ-potential of unsized vesicles showed a slight variation in parabolic pattern with a minimum at the centre of the variable concentrations in the organic-phase. Maximum PDI 0.579 and minimum ζ-potential −20.4 mV were observed at the highest dilution of HPCE/CHCl_3_ 7.5 μM/ml using distilled water as dilution medium. The variable Z-av and ζ-potential values at variable HPCE/CHCl_3_ concentration showed a direct relationship (Table-3). PDI of vesicles after sizing was significantly improved in the range of 0.15 to 0.095. The ζ-potential of vesicles was slightly increased but more uniform after sizing in the range of −18.15 mV to −17.1 mV basing size-dependent variation.

#### 3.4.2 Atomic force microscope topography

Liposomal stock dispersion after 100x dilution in DW was imaged through atomic force microscopy in amplitude-detection mode (AC-AFM). The stock dispersion contained HPCE/DW 4 μmol/ml and Cho 55.6 mol% of total solids. Liposomes were observed within 5-10 min of deposition on silicon-wafers. The promptness in observation was critical to minimize topographical changes due to substrate-sample interaction and or environmental effects [39]. Spheroidal vesicles having size-dependent surface-homogeneity signified aptness of the sample preparation method and the scanning technique used (Figure-2). However, a probable stretching effect of the cantilever, sample-substrate interaction and flaccidity of the liposomes lent a difference in observed length to width ratio (Figure-2b & e) [40]. The vesicles maintained their integrity due to lamellar elasticity letting a slight bent during the scanning process. The observed diameter of the sized vesicles was in accordance with the pore size of the smallest filter-membrane (Table-3). The height of samples was accorded to the size and rigidity of samples. A little difference in the height of sized samples to corresponding unsized vesicles was observed. Mean diameter, height, average surface roughness (Ra) and rigidity were improved by reducing the size of any given sample (Figure-2, Table-3). Surface homogeneity was confirmed from 3D topographic images analysed in the process mode of an inbuilt software (WINSPM 5). Section view and three-dimensional imaging plot indicated a smooth and regular surface profile of the observed liposomes. Surfaces of sized vesicles appeared more uniform than the corresponding unsized samples (Figure-2c & f). Invitro rigidity of liposomes was assessed from the height data of individual vesicles through AC-AFM in combination of Z-av measured after 100x dilution in 10 μM NaCl. The data of Z-av in saline medium resembled the length of vesicles measured through AFM, hence, considered suitable to asses rigidity. Comparing various concentrations of HPCE/CHCl_3_, 15 μmol/ml sized samples showed maximum rigidity of 0.14±0.03, height 30.58±6.82 nm, Z-av 149.25±30.2 nm, and roughness 5.15±1.68 nm. The calculated volume was size-dependent and progressively increasing with the decreasing molar concentration of amphiphiles in the organic-phase. However, more uniformity in volumes was observed at HPCE/CHCl_3_ 15μmol/ml of unsized vesicles. Majority of the liposomes appeared intact and spheroidal in the obtained images. However, slightly flattened structures were observed due to the flaccid nature of liposomes obvious from their length to height ratio (Table-3 & Figure-2).

**Figure-2:**
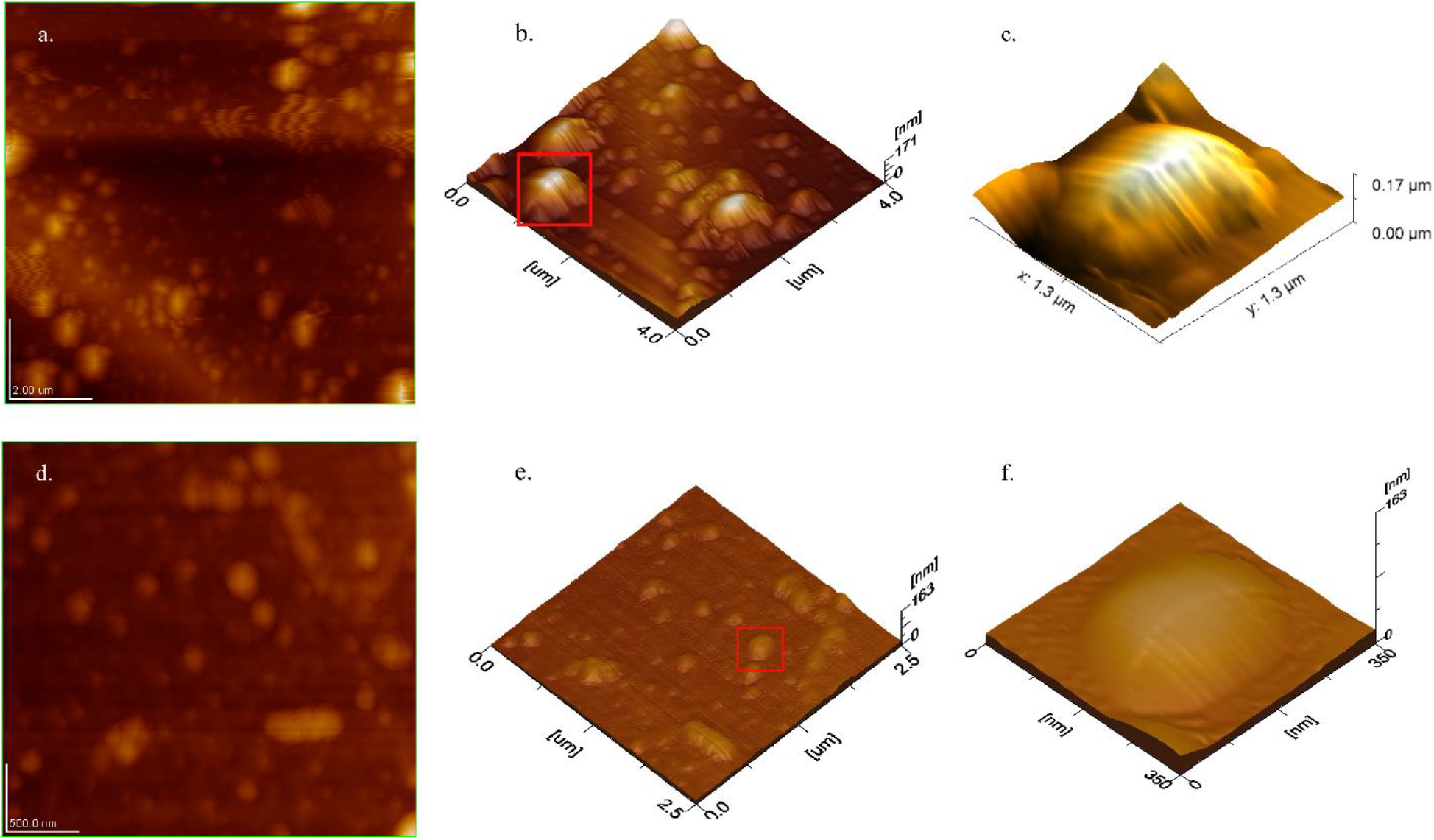
AFM topography of liposomes produced with optimized parameters showing comparison between sized and unsized liposomes **a:** 2D image of unsized liposomes **b:** 3D image of the same sample **c:** 3D image of selected single liposomes showing larger size, more height and surface roughness **d:** 2D image of sized liposomes **e:** 3D image of sized liposomes **f:** 3D image of selected single liposomes showing flattened and smaller size and size-dependent lower height but smooth surface and high rigidity.

#### 3.4.3 Entrapped aqueous volume

High entrapped aqueous volumes were observed consistent with the calculated volumes (Table-3). A slight variation in variable HPCE/CHCl_3_ ranged from 20.09±0.18 to 23.87±0.18 L/mol also reflected size dependence. Unsized liposomes were excluded from the study due to broader PDI and hence of little practical implication. However, the long dialysis period of 24hrs may have some leakage effects which were not included in the calculations. So, the given values might indicate a minimum achievable entrapped volume of the available aqueous phase.

#### 3.4.4 Accelerated stability studies

The liposomal suspension remained uniform in type-1 glass vials throughout the study. However visible precipitates/aggregates appeared in the plastic falcon tubes for the same set of experiments and hence excluded from the study. During the first month of storage at both temperature conditions in type-1 glass vials, a slight steep both in PDI and Z-av was observed (Figure-3d). However, the values of PDI gradually declined but Z-av increased over an extended period of storage showing aggregation of vesicles. Samples stored in refrigerator (REF) showed a minute increase in size from 226.3 nm to 246.6 nm and PDI from 0.105 to 0.185 (Figure-3a & b) over a period of 6 months. However, comparatively bigger change was observed at room temperature (RT) where Z-av raised from 226.3 nm to 327.5 nm and PDI 0.105 to 0.214 (Figure: 3a & c).

**Figure 3.**
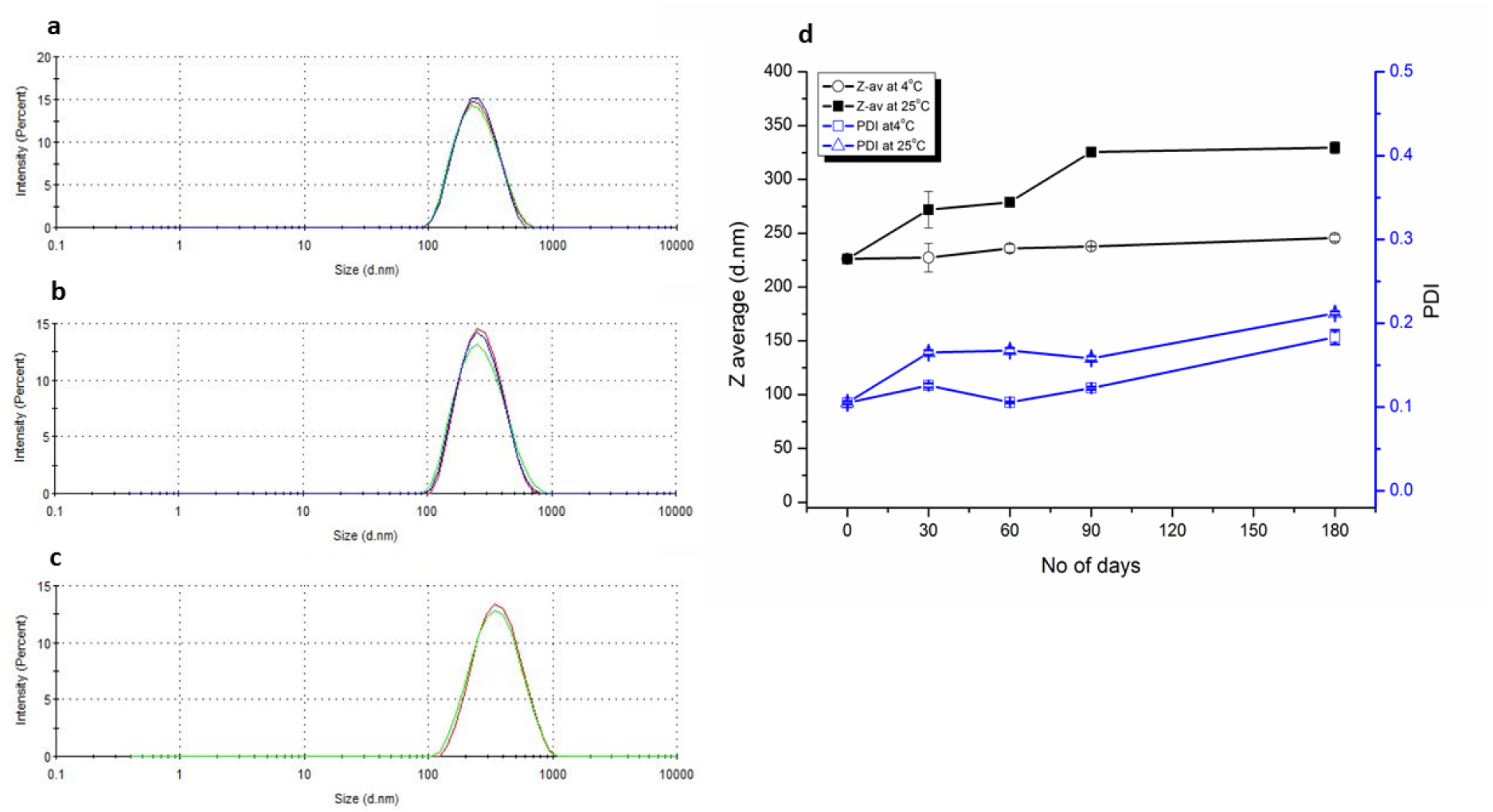
Accelerated stability study of optimized liposomes after sizing by serial membrane filtration with 0.2 μm membrane filter and stored in the primary hydration medium ^**a:**^ DLS results at 0-day ^**b:**^ DLS results after 180 days at 4°C ^**c:**^ DLS results after 180 days at 25°C ^**d:**^ Z-av and PDI vs number of days at 4°C and 25°C, the results are presented as mean ± SD

#### 3.4.5 Process validation

The batch sizes of optimized formulations were scaled-up by 5x increment and evaluated for reproducible results. A variety of phospholipids based on their hydrogenation and source material were used to prepare liposomes under an optimized set of parameters and checked for reproducible results (data not included in this paper). All formulations were produced by using optimized formulation and processing parameters (Section-3.2). The three reproducible trial formulations and 5x scaled-up batches yielded comparable results in terms of size and PDI.

## 4 Discussion

### 4.1 Key features of optimization studies

The experimentation was aimed to use higher solids/solvent ratio at possibly lower temperature imperative to high encapsulation efficiency particularly of thermosensitive molecules [4]. Samples produced at above and below 45°C showed precipitation or aggregation while using high solid/solvent ratio. Z-av of optimized formulations was gradually decreasing with declining molar concentration of HPCE/CHCl_3_ μmol/ml and vice versa [23, [41] against fixed aqueous volume. It presumed the slow movement of diluted phospholipids from organic to aqueous phase favouring the formation of smaller sized vesicles. Whilst the PDI was slightly parabolic from high to lower HPCE/CHCl_3_ μmol/ml. Suggesting DW/CHCl_3_ as 4:1 or HPCE/CHCl_3_ 15 μmol/ml was a suitable concentration for comparable smaller size and PDI of liposomes. It suggested that lower temperatures and concentrated amphiphiles in the organic-phase collectively resulted in condensation and tactless relocation into the aqueous phase. While at higher temperatures (55°C) although favoured faster removal of organic phase but tendered the solid components in straightened molecular dimension or GUVs and consequent aggregation. The concentration of Cho was found critical for vesicle formation with an optimal level of HEPC/Cho as a 4:5 molar ratio. The concentration below this level promoted vesicle aggregation if not sized while the above levels favoured the precipitation of Cho. The sizing of liposomes by serial membrane filtration is associated with concomitant deformation and reformation of vesicles during extrusion through narrow orifices of the filter membrane. Therefore, the extrusion of LUVs in the primary hydration medium is essential for the restoration of dissolved components [35].

### 4.2 Properties of optimized CI and SPT liposomes

By evading injection phase of CI-method resulted in the SPT-method without any significant variation in the quality of liposomes but furnished process efficiency (Figure-1a). Therefore, SPT-method may be preferred for its further simplicity, handiness, least interaction of organic solvent to aqueous components and better yield. A high solid solvent ratio signified high encapsulation efficiency of aqueous drugs [4]. While immiscible organic solvent gave true molecular dispersion of organic components ensuing good entrapment of lipophiles in the lamellar compartment. Attempts were made to analyse the residual traces of CHCl_3_ in the final product but couldn’t obtained reproducible results after 10 minutes of rotary evaporation under vacuum. However its complete removal is evidenced from the previous reports [24, [27] and overall volume loss of total aqueous suspension (Section-3.4). The desired size distribution required essential steps of sizing [34]. The CI liposomes can be produced over a wide range of temperatures (25-45 °C) but require large aqueous volumes at low temperatures signifying low encapsulation efficiency. The liposomal size gradually increased with increasing phospholipids concentration in both or any solvent phase [5, [29]. The aggregation of vesicles at 55 °C (hydrated transition temperature of HPCE) infers the formation of larger globules due to the high fluidity of phospholipid molecules. Cho 55.6 mol% yielded uniform vesicles while lower concentrations indicated phase segregation [42, [43]. High Cho contents not only aided the shelf-life stability of liposomes but efficient encapsulation of aqueous drugs as well [32, [33].

Size and surface charge on liposomes are imperative in the stability, kinetics, biodistribution, enhanced permeation and retention, sterilization and interaction with the targeted cells [44]. Usually, the liposomes are sized by membrane filtration at above the phase transition temperature of lipids [34]. The pore size of the membrane used for extrusion determines the size of vesicles with more uniformity compare to other methods [7, [35]. In this study, the liposomal size was effectively reduced at the processing temperature that was ≤10°C below the transition temperature of lipid in use. Observing above mentioned formulations in 10 μM NaCl by DLS and topographical images of NC-AFM (Table-3, Figure-2) indicated electrical polarity dependant size distribution in DW. The type of hydration medium for the production of vesicles play a crucial role in size distribution where DW lent lower size and broader PDI [45]. The aim of using DW as a hydration medium in this study was to investigate various parameters in the very raw form to leave a room for improvement while working with drug-loaded formulations.

AFM is the most suitable non-invasive technique to observe the liposomes and soft biological samples without staining and under atmospheric conditions [36, [46] based on its simplicity and high resolution approaching 1Å [39]. While both contact and non-contact AFM have the drawbacks of damaging soft vesicles and low resolution respectively [36]. AC-AFM exerts low pressure while operating at separation of 5-20Å and high resolution by intermittent tapping. Variation in the height to width ratio of AC-AFM images depicted size-dependent rigidity and flaccid nature of the vesicles [37]. Being anionic vesicles deposited on the silicon wafers with a slight negative surface-charge reduced the sample-substrate interaction [47]. The diameter measured by AC-AFM was smaller than the hydrodynamic diameter through DLS and in accordance with the smallest filter-membrane used for sizing. The larger Z-av diameter was electrical potential dependent with ζ-potential −17.3±0.6 mV. Other electron microscopy techniques require staining of samples and observations under high vacuum which affect the delicate structure of liposomes. The coating process for SEM topography collapsed the flaccid liposomes from globular to circular objects. The unilamellar images having sharp boundaries were slightly larger in diameter than the AFM images. This method gives gross estimation of lamellarity by staining interlamellar spaces and resolve the lamellar walls [48, [49]. Rigidity is an important factor aiding to the stability of liposomes particularly if a long systemic circulation times is required. High Cho contents, nature of phospholipids, surface potential and coating can aid to the rigidity of a given size of liposomes [37].

High encapsulated concentration is beneficial to reduce the dose size, increased dosing interval and make the product cost-effective. Ideally for a 130 nm liposomes entrapped volume will be 36 L/mol, by ether injection it was reported 14+6L/mol. [19] HPCE/Cho 50 mol% gave 16.41 L/mol and filtered 0.22 μm filter gave 9.0 L/mol. [22, [23] In current methods, the de-novo assemblage and high solid/solvent ratio in aqueous phase favoured large entrapped volume (Table-3) ensuing high encapsulation of hydrophiles. [50]

### 4.3 Vesiculation pathway

The HPCE/Cho mixture yielded a true solution in CHCl_3_. In the CI-method the amphiphiles spontaneously reoriented in the aqueous-phase as leaving orifice of the needle. The polar head interacted with water molecules while nonpolar tails sequestered in bilayer formed by the adjacent chains of amphiphiles [51]. The resultant highly unstable milky suspension was a mixture of vesicles and micelles containing both solvents. Removal of immiscible organic solvent stabilized the lamellar globules by holding water on both sides. However, for concentrated solid/solvent dispersions a minimum temperature of 45°C helped to prevent precipitation before assemblage into the lamellar vesicles.

In SPT method organic-phase containing amphiphiles was partitioned at the bottom of the flask. CHCl_3_ readily exchanged solute molecules while moving through the aqueous-phase during rotary evaporation. Amphiphiles reoriented on contact with water to minimize surface free energy by sequestering hydrophobic tails in the bilayers [51]. The size of the vesicles was a function of fragmentation achieved through swirling of the aqueous phase and the rate of evaporation of organic solvent to exchange phospholipids. This phenomenon was evidenced by larger size vesicles due to gradient solid/solvent ratio in organic-phase (Table-3).

### 4.4 Stability of the liposomes

Both methods suddenly relocate HPCE/Cho indicating uniform lamellar distribution [27] aiding to the stability of liposomes. Cho enhances the strength of liposomes by modifying fluidity and interaction of phospholipids in the lamellar membranes [13, [39]. The saturated acyl chain of HPCE has strong interaction with the steroidal ring of Cho molecule. Hence Cho concentration (55.6 mol%) showed high compressibility and low permeability to water [1, [33] during storage. Therefore, the sole reason for stability might be a high molar concentration of Cho [37]. Storage at REF restricted molecular movement and hence stronger lamellae [14]. A size-dependent ζ-potential −17.1+0.6 mV also aided in the stability of formulation by exhibiting inter-vesicular repulsion [5]. A slight increase in the size of liposome at both temperatures indicated swelling of the bilayers while stored in the primary hydration medium. Overwhelmingly the shelf-life stability can be improved by further enhancing the ζ-potential and lyophilization with suitable cryoprotectant [4, [5].

### 4.5 Scalability

The injection phase of the CI-method may restrict the method for small batches until some mechanical device for injection is adapted. However, the SPT-method was more flexible to be scaled up and down due to the simple manufacturing protocol and equipment required for production. 5x scale-up studies showed reproducible results from 12 ml to 300 ml batch sizes. A maximum of half of the rotating flask of the rotary evaporator may be occupied to allow sufficient surface area for evaporation.

### 4.6 Pros and cons of the methods

The methods described in this paper were advantageous in terms of swiftness, reproducibility, productivity, adaptability, homogeneity and large entrapped volume. The process loss of CI-method was effectively compensated in the SPT-method where the whole material was subjected to vesiculation. However, comparatively lower size and PDI were the overriding features of CI over SPT-method. The use of immiscible organic solvent and vacuum evaporation facilitated easy removal and minimum interaction with aqueous components benefiting sensitive molecules. Use of lower temperature i.e. ≤10°C below the transition temperature of lipids and short processing time will aid in the encapsulation of thermosensitive drugs. Processing at 25°C was possible in large aqueous volume but loss of diluted solutes and hence low encapsulation efficiency is obvious. The liposomes may be used fresh or stored in light protected type-1 glass in the liquid form for a longer period at 5±3°C. However, the potential disadvantages include cumbersome size reduction by serial membrane filtration for the smaller size and PDI. The unentrapped material if not desired must be removed by washing through dialysis at 5±3°C to avoid aggregation and size growth of the vesicles.

### 4.7 Comparison with other methods

The current development aimed a simple method in terms of required equipment, processing time, versatility and handiness while preparing high-quality liposomes. Unlike film hydration methods [2, [4, [12] intermediate solid phase formation was bypassed ensuing compositionally homogeneous and stable lamellae besides using high molar concentrations of Cho (55.6 mol%). Large entrapped volume than film hydration methods was ascription of spontaneous rearrangement of phospholipids. Swiftly sizing by membrane filtration of CI and SPT liposomes within the same hydration medium was advantageous upon other size-reduction methods like sonication and homogenization [35] in retaining the entrapped entities.

Bulk methods give comparable results to film hydration methods in terms of quality but have several limitations. Large aqueous volumes were used in majority methods leading to high entrapped volume but low encapsulation efficiency [19, [20]. Contrariwise CI and SPT methods used concentrated solutions and bulk methods to overcome the aforementioned limitations. Similarly, the use of miscible organic solvents and or high temperature [19, [20, [21, [24, [26] limited some methods to the stable drugs under such conditions and risk of leakiness. Exclusion of Cho [20, [24] certainly favoured vesiculation at lower temperatures but led a compromised physical stability and lower entrapment efficiency [33]. Alternatively, CI and SPT methods used high Cho contents in immiscible organic solvent and temperature ≤10°C below the transition temperature of the lipid employed. CI-method was applicable at lower temperatures (25°C) but higher aqueous volumes limited its suitability due to low encapsulation of diluted hydrophiles. Even though some methods employed immiscible organic solvents, high lipid concentrations but highly sophisticated equipment and or procedures which confine their commonness [23, [27, [28, [29, [30, [31]. Instead, the CI-method needed an injection system followed by rotary evaporation but the SPT-method even so, eliminated the injection step from the process.

## Conclusion

Collectively both CI and SPT methods tendered broader solution to variety of parameters for the preparation of liposomes like composition, temperature, Z-av, and scalability. Simple technique and equipment were used to obtain the desired size distribution. A processing temperature 10°C below the transition temperature of different phospholipids with 55.6 mol percent Cho was effective in formulating high solid solvent ratio like HPCE 4 μmol/ml of aqueous phase. Processing temperature was lowered further by using dilute solutions of solids in any or both phases facilitating encapsulation of thermophiles. However minimum possible volume of organic phase was always preferred for its effective removal to safeguard shelf life and quality of entrapped entities. High solid/solvent ratio resulted in larger size vesicles but imperative to higher entrapment efficiency of hydrophiles. Therefore, sizing at processing temperature in primary hydration medium was pivotal to retain the initial concentration of entrapped molecules while attaining desired Z-av and PDI. The produced liposomes having high Cho contents were stable as per ICH guidelines at REF while packed in light protected type-1 glass containers. However, lyophilization with suitable cryoprotectant depending upon the nature of phospholipids and entrapped molecules will definitely enhance shelf life quality of the product. The use of immiscible organic phase assisted to entrap both lipophilic and hydrophilic molecules simultaneously signifying its equal appliance in diversified fields using nano formulations. Validation and scale-up studies ensued the handiness of SPT method for a variety of amphiphiles and scaleup for commercial production or down to point of care use. However, the CI method is currently restricted to small batches only until a suitable mechanical equipment is adapted. Entrapment of various lipophilic and hydrophilic molecules and their stability in the intended dosage-forms need to be evaluated. Once the simple equipment is set up the LUVs can be obtained within 20 mins followed by sizing if needed.

## Data Availability Statement

All data generated and analysed during the current study are included in this published article. Further details and support are available from corresponding author on reasonable request.

## Authors Contribution Statement

**MIK:** Conceptualization, Investigation, Data Curation, Writing-Original Draft

**NA:** Methodology, Validation

**MFU:** Software, Visualization

**AR:** Formal analysis

**NMA:** Resources

**GMK:** Supervision, Project administration, Writing-Review and Editing.

## Referencess

1. Crosasso P, Ceruti M, Brusa P, Arpicco S, Dosio F, Cattel L. Preparation, characterization and properties of sterically stabilized paclitaxel-containing liposomes. Journal of Controlled Release. 63(1-2), 19–30 (2000).

2. Shigehiro T, Kasai T, Murakami M, Sekhar SC, Tominaga Y, Okada M, Kudoh T, Mizutani A, Murakami H, Salomon DS. Efficient drug delivery of paclitaxel glycoside: A novel solubility gradient encapsulation into liposomes coupled with immunoliposomes preparation. PloS one. 9(9), e107976 (2014).

3. Patil YP, Jadhav S. Novel methods for liposome preparation. Chemistry and physics of lipids. 177(8-18 (2014).

4. Maione-Silva L, de Castro EG, Nascimento TL, Cintra ER, Moreira LC, Cintra BAS, Valadares MC, Lima EM. Ascorbic acid encapsulated into negatively charged liposomes exhibits increased skin permeation, retention and enhances collagen synthesis by fibroblasts. Scientific reports. 9(1), 522 (2019).

5. Chorachoo J, Amnuaikit T, Voravuthikunchai SP. Liposomal encapsulated rhodomyrtone: A novel antiacne drug. Evidence-Based Complementary and Alternative Medicine. 2013((2013).

6. Vyas S, Kannan M, Jain S, Mishra V, Singh P. Design of liposomal aerosols for improved delivery of rifampicin to alveolar macrophages. International journal of pharmaceutics. 269(1), 37–49 (2004).

7. Dar MJ, McElroy CA, Khan MI, Satoskar AR, Khan GM. Development and evaluation of novel miltefosine-polyphenol co-loaded second generation nano-transfersomes for the topical treatment of cutaneous leishmaniasis. Expert Opinion on Drug Delivery. 1–14 (2019).

8. Lasic D. Liposomes. American Scientist. 80(1), 20–31 (1992).

9. Torchilin V. Tumor delivery of macromolecular drugs based on the epr effect. Advanced drug delivery reviews. 63(3), 131–135 (2011).

10. Ying X, Wen H, Lu W-L, Du J, Guo J, Tian W, Men Y, Zhang Y, Li R-J, Yang T-Y. Dual-targeting daunorubicin liposomes improve the therapeutic efficacy of brain glioma in animals. Journal of Controlled Release. 141(2), 183–192 (2010).

11. Wang J, Kang Y-X, Pan W, Lei W, Feng B, Wang X-J. Enhancement of anti-inflammatory activity of curcumin using phosphatidylserine-containing nanoparticles in cultured macrophages. International journal of molecular sciences. 17(6), 969 (2016).

12. Le U, Ngo D, Nguyen T, Nguyen Q, Ton J. Enhanced selective cytotoxicity in pancreatic cancer cells using egf-conjugated liposome-encapsulated curcumin. International Conference on the Development of Biomedical Engineering in Vietnam. 217–221 (2017)

13. Mugabe C, Azghani AO, Omri A. Liposome-mediated gentamicin delivery: Development and activity against resistant strains of pseudomonas aeruginosa isolated from cystic fibrosis patients. Journal of Antimicrobial Chemotherapy. 55(2), 269–271 (2005).

14. Huang W-C, Deng B, Lin C, Carter KA, Geng J, Razi A, He X, Chitgupi U, Federizon J, Sun B. A malaria vaccine adjuvant based on recombinant antigen binding to liposomes. Nature nanotechnology. 13(12), 1174 (2018).

15. Le UM, Cui Z. Biodistribution and tumor-accumulation of gadolinium (gd) encapsulated in long-circulating liposomes in tumor-bearing mice for potential neutron capture therapy. International journal of pharmaceutics. 320(1-2), 96–103 (2006).

16. Bally M, Bailey K, Sugihara K, Grieshaber D, Vörös J, Städler B. Liposome and lipid bilayer arrays towards biosensing applications. Small. 6(22), 2481–2497 (2010).

17. Wagner A, Vorauer-Uhl K. Liposome technology for industrial purposes. Journal of Drug Delivery. 2011((2011). doi:10.1155/2011/591325

18. Szoka F, Papahadjopoulos D. Procedure for preparation of liposomes with large internal aqueous space and high capture by reverse-phase evaporation. Proceedings of the national academy of sciences. 75(9), 4194–4198 (1978).

19. Deamer D, Bangham A. Large volume liposomes by an ether vaporization method. Biochimica et Biophysica Acta (BBA)-Nucleic Acids and Protein Synthesis. 443(3), 629–634 (1976).

20. Batzri S, Korn ED. Single bilayer liposomes prepared without sonication. Biochimica et Biophysica Acta (BBA)-Biomembranes. 298(4), 1015–1019 (1973).

21. Ishii F, Takamura A, Ishigami Y. Procedure for preparation of lipid vesicles (liposomes) using the coacervation (phase separation) technique. Langmuir. 11(2), 483–486 (1995).

22. Deamer DW. Preparation and properties of ether-injection liposomes. Annals of the New York Academy of Sciences. 308(1), 250–258 (1978).

23. Kim S, Jacobs RE, White SH. Preparation of multilamellar vesicles of defined size-distribution by solvent-spherule evaporation. Biochimica et Biophysica Acta (BBA)-Biomembranes. 812(3), 793–801 (1985).

24. Moscho A, Orwar O, Chiu DT, Modi BP, Zare RN. Rapid preparation of giant unilamellar vesicles. Proceedings of the National Academy of Sciences. 93(21), 11443–11447 (1996).

25. Kagawa Y, Racker E. Partial resolution of the enzymes catalyzing oxidative phosphorylation xxv. Reconstitution of vesicles catalyzing 32pi—adenosine triphosphate exchange. Journal of Biological Chemistry. 246(17), 5477–5487 (1971).

26. Oku N, Scheerer JF, MacDONALD RC. Preparation of giant liposomes. Biochimica et Biophysica Acta (BBA)-Biomembranes. 692(3), 384–388 (1982).

27. Buboltz JT, Feigenson GW. A novel strategy for the preparation of liposomes: Rapid solvent exchange. Biochimica et Biophysica Acta (BBA)-Biomembranes. 1417(2), 232–245 (1999).

28. Kuroiwa T, Kiuchi H, Noda K, Kobayashi I, Nakajima M, Uemura K, Sato S, Mukataka S, Ichikawa S. Controlled preparation of giant vesicles from uniform water droplets obtained by microchannel emulsification with bilayer-forming lipids as emulsifiers. Microfluidics and nanofluidics. 6(6), 811 (2009).

29. Mijajlovic M, Wright D, Zivkovic V, Bi J, Biggs MJ. Microfluidic hydrodynamic focusing based synthesis of popc liposomes for model biological systems. Colloids and Surfaces B: Biointerfaces. 104(276–281 (2013).

30. Castor TP. Methods and apparatus for making liposomes. (1996)

31. Akashi K-i, Miyata H, Itoh H, Kinosita Jr K. Preparation of giant liposomes in physiological conditions and their characterization under an optical microscope. Biophysical journal. 71(6), 3242–3250 (1996).

32. Grant GJ, Barenholz Y, Piskoun B, Bansinath M, Turndorf H, Bolotin EM. Drv liposomal bupivacaine: Preparation, characterization, and in vivo evaluation in mice. Pharmaceutical research. 18(3), 336–343 (2001).

33. de Meyer F, Smit B. Effect of cholesterol on the structure of a phospholipid bilayer. Proceedings of the National Academy of Sciences. 106(10), 3654–3658 (2009).

34. Olson F, Hunt C, Szoka F, Vail W, Papahadjopoulos D. Preparation of liposomes of defined size distribution by extrusion through polycarbonate membranes. Biochimica et Biophysica Acta (BBA)-Biomembranes. 557(1), 9–23 (1979).

35. Meisel JW, Gokel GW. A simplified direct lipid mixing lipoplex preparation: Comparison of liposomal-, dimethylsulfoxide-, and ethanol-based methods. Scientific reports. 6(27662 (2016).

36. De Oliveira R, Albuquerque D, Cruz T, Yamaji F, Leite F. Measurement of the nanoscale roughness by atomic force microscopy: Basic principles and applications. Atomic force microscopy-imaging, measuring and manipulating surfaces at the atomic scale. 147–175 (2012).

37. Nakano K, Tozuka Y, Yamamoto H, Kawashima Y, Takeuchi H. A novel method for measuring rigidity of submicron-size liposomes with atomic force microscopy. International journal of pharmaceutics. 355(1-2), 203–209 (2008).

38. Guideline IHT. Stability testing of new drug substances and products. Q1A (R2), current step. 4(1-24 (2003).

39. Ruozi B, Tosi G, Forni F, Fresta M, Vandelli MA. Atomic force microscopy and photon correlation spectroscopy: Two techniques for rapid characterization of liposomes. European Journal of Pharmaceutical Sciences. 25(1), 81–89 (2005).

40. Bolean M, Borin IA, Simão AM, Bottini M, Bagatolli LA, Hoylaerts MF, Millán JL, Ciancaglini P. Topographic analysis by atomic force microscopy of proteoliposomes matrix vesicle mimetics harboring tnap and anxa5. Biochimica et Biophysica Acta (BBA)-Biomembranes. 1859(10), 1911–1920 (2017).

41. Pons M, Foradada M, Estelrich J. Liposomes obtained by the ethanol injection method. International journal of pharmaceutics. 95(1-3), 51–56 (1993).

42. Redondo-Morata L, Giannotti MI, Sanz F. Influence of cholesterol on the phase transition of lipid bilayers: A temperature-controlled force spectroscopy study. Langmuir. 28(35), 12851–12860 (2012).

43. Maestrelli F, González-Rodríguez ML, Rabasco AM, Mura P. Preparation and characterisation of liposomes encapsulating ketoprofen–cyclodextrin complexes for transdermal drug delivery. International journal of pharmaceutics. 298(1), 55–67 (2005).

44. Porfire A, Muntean DM, Rus L, Sylvester B, Tomuţă I. A quality by design approach for the development of lyophilized liposomes with simvastatin. Saudi Pharmaceutical Journal. 25(7), 981–992 (2017).

45. Obeid MA, Khadra I, Mullen AB, Tate RJ, Ferro VA. The effects of hydration media on the characteristics of non-ionic surfactant vesicles (nisv) prepared by microfluidics. International journal of pharmaceutics. 516(1-2), 52–60 (2017). doi:https://doi.org/10.1016/j.ijpharm.2016.11.015

46. Ruozi B, Belletti D, Tombesi A, Tosi G, Bondioli L, Forni F, Vandelli MA. Afm, esem, tem, and clsm in liposomal characterization: A comparative study. International journal of nanomedicine. 6(557 (2011).

47. Palmer AF, Wingert P, Nickels J. Atomic force microscopy and light scattering of small unilamellar actin-containing liposomes. Biophysical journal. 85(2), 1233–1247 (2003).

48. Perkins W, Minchey S, Ahl P, Janoff A. The determination of liposome captured volume. Chemistry and physics of lipids. 64(1-3), 197–217 (1993).

49. Miyamoto VK, Stoeckenius W. Preparation and characteristics of lipid vesicles. The Journal of membrane biology. 4(1), 252–269 (1971).

50. Xu X, Khan MA, Burgess DJ. A quality by design (qbd) case study on liposomes containing hydrophilic api: Ii. Screening of critical variables, and establishment of design space at laboratory scale. International journal of pharmaceutics. 423(2), 543–553 (2012).

51. Lasic D. Mechanisms of liposome formation. Journal of liposome research. 5(3), 431–441 (1995).

